# Herbicide metolachlor alters gene expression and reshapes the interaction between a bloom-forming cyanobacterium and its chytrid parasite

**DOI:** 10.1101/2025.10.07.680865

**Authors:** Alice Balard, Jürgen F. H. Strassert, Justyna Wolinska, Erika B. Martínez-Ruiz

## Abstract

Metolachlor (MET) is among the most widely used herbicides worldwide and a common freshwater contaminant. Beyond direct toxicity, MET can reshape ecological interactions, including those between bloom-forming cyanobacteria and their chytrid parasites. Although MET is known to reduce chytrid fitness and alter cyanobacterium–chytrid interaction, the molecular basis of these effects remains unknown. Using dual RNA-sequencing, we analysed the gene expression of the toxigenic cyanobacterium *Planktothrix agardhii* and its obligate chytrid parasite *Rhizophydium megarrhizum* under exposure to an environmentally relevant concentration of MET. MET reprogrammed the parasite *R. megarrhizum* by altering the expression of genes governing developmental transitions, xenobiotic stress responses and intracellular signalling in both free-living and infecting cells. The transcriptional signature that typically distinguishes zoospores from infecting chytrids disappeared under MET exposure. This was not due to an arrest of the infection process, but rather to an increase in the baseline expression of infection-associated genes in zoospores, thus blurring the dispersal-to-infection transition. In contrast, *P. agardhii* showed no transcriptional response to MET in the absence of infection, whereas co-exposure to both the parasite and MET was associated with transcriptional changes related to energy acquisition and carbohydrate metabolism. Overall, our results suggest that MET weakens chytrid top-down control and favours cyanobacterial persistence, with consequences for bloom formation and energy transfer through aquatic trophic webs. Our study provides the first replicated molecular view, using a dual-RNA transcriptomic approach, of how herbicide exposure reshapes the interaction between a bloom-forming cyanobacterium and its chytrid parasite.

## Introduction

Cyanobacteria are primary producers that form the base of most aquatic trophic webs, along with other phytoplankton [1]. In response to eutrophication and other global environmental changes, cyanobacteria often form blooms [2, 3]. These blooms can deplete oxygen and produce cyanotoxins, which are potentially harmful compounds that pose an ecological and health risk [4, 5]. Cyanobacterial growth can be regulated by fungal parasites from the phylum Chytridiomycota, known as chytrids [6]. The chytrid life cycle begins with a free-living zoospore that attaches to a host cell, develops rhizoids to extract nutrients and forms a sporangium from which new zoospores are released, resulting in lethal consequences for the host [7]. Through this process, chytrids influence bloom dynamics, nutrient cycling [6, 8, 9] and cyanobacterial diversity [10, 11]. Both chytrids and phytoplankton, including cyanobacteria, play key roles in aquatic ecosystems, but their interactions are vulnerable to anthropogenic pollutants. For instance, exposure to various pollutants can reduce chytrid parasite fitness [12–16], while pollutant effects on hosts may depend on parasite presence [17]. For example, diclofenac and cigarette butt leachate inhibit host growth in the absence of chytrids, but can enhance it when chytrids are present by reducing parasitic pressure [13, 16].

Herbicides are the most widely used group of pesticides worldwide, accounting for roughly half of total global pesticide use [18]. Metolachlor (MET), a chloroacetanilide herbicide, ranks among the most commonly used herbicides worldwide [19]. Although MET concentrations in surface waters usually fall within the ng L^-1^ to low μg L^-1^ range [20, 21], substantially higher levels have also been reported [22, 23]. MET can affect aquatic organisms, including cyanobacteria and chytrids, and may modify their interaction. Previous work in this system showed that short-term MET exposure promoted cyanobacterial growth in the absence of chytrids but not during co-exposure, whereas long-term exposure reduced chytrid fitness [14]. A complementary multi-biomarker study showed metabolic stress in cyanobacteria under co-exposure, but not under MET exposure alone [24]. The molecular basis of these effects remains unknown.

Building on previous data on growth, fitness and biomarkers, this study provides molecular insights into how MET affects cyanobacteria, chytrids and their interaction, with implications for bloom dynamics and energy transfer along the trophic web in polluted environments. We therefore hypothesised that MET 1) alters the expression of genes associated with host colonisation and nutrient uptake in chytrid zoospores and 2) promotes the overexpression of oxidative stress-related genes in infected but not uninfected cyanobacteria.

## Materials and Methods

### Cyanobacteria and chytrids culturing conditions

The host–parasite system consisted of the toxigenic filamentous cyanobacterium *Planktothrix agardhii* strain NIVA-CYA630 and its obligate chytrid parasite *Rhizophydium megarrhizum* strain Chy-Kol2008 [25]. Cyanobacteria were routinely maintained in Z8 medium [26] at 16 °C, under a continuous light intensity of 20 µmol photons m^-2^ s^-1^. Chytrids were cultivated by transferring zoospores to uninfected cyanobacterial cultures every three weeks. Both organisms have been maintained under continuous light for more than a decade and are therefore acclimated to this light regime.

### Metolachlor (MET) exposure

Infected and uninfected cyanobacterial cultures were exposed to 100 μg L^-1^ MET, an environmentally relevant concentration previously shown to affect chytrid fitness [14]. The experiment included 20 units with two MET treatments (unexposed vs. exposed), two infection conditions (uninfected vs. infected) and five replicates each (**Figure 1**; **Table S1**).

**Figure 1.**
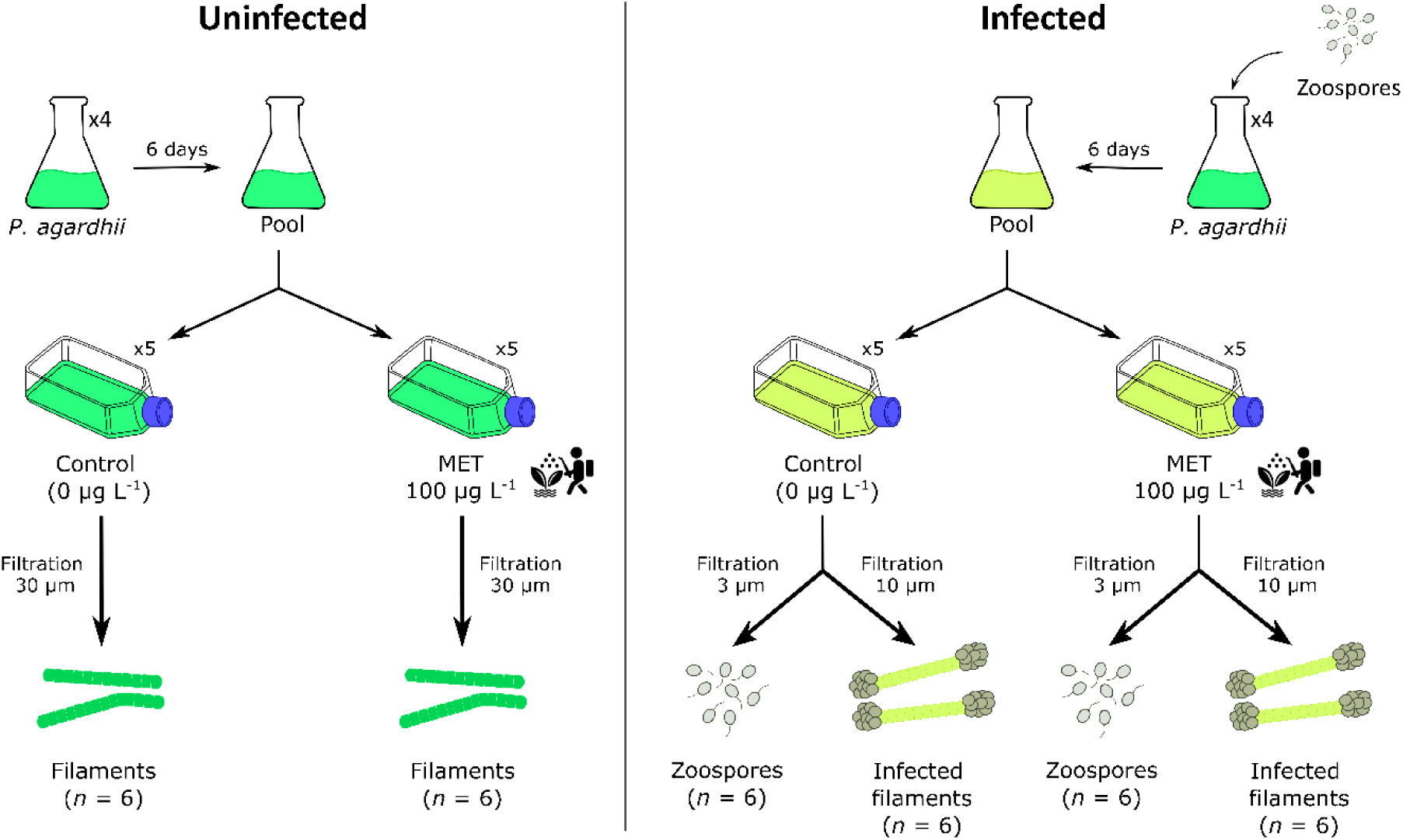
Scheme of the experimental design. Different number of cells were collected for each sample type: 30–34 uninfected cyanobacterial filaments, 40–44 free-living chytrid zoospores and 6–11 infected cyanobacterial filaments, each containing approximately 10 encysted zoospores or sporangia. The number of samples per treatment is represented by *n*. As one replicate from each treatment was sampled twice as a technical replicate, six samples were obtained for each sample type.

To produce chytrid zoospores for infection, an exponentially growing cyanobacterial culture was infected 10 days before the experiment. Afterwards, the culture was filtered through a 5 μm nylon mesh and a 3 μm polycarbonate filter to obtain the zoospore suspension [10]. Zoospore density was quantified using a Sedgewick-Rafter chamber under an inverted microscope (Nikon Ti Eclypse).

During the two weeks prior to MET exposure, eight cyanobacterial cultures were maintained as exponentially growing semi-continuous cultures by dilution with sterile Z8 medium to an OD750nm of 0.05 thrice per week. Four cultures were inoculated with 750 zoospores mL^-1^ and incubated for six days to allow the infection to establish; the remaining four uninfected cultures followed the same incubation schedule. Infection was confirmed by quantifying infection prevalence (i.e. the proportion of filaments containing encysted zoospores or sporangia) under an inverted microscope. On the day of MET exposure, uninfected cultures were diluted to an OD750nm of 0.05, whereas infected cultures were not diluted to avoid disrupting host–parasite interactions. Both culture types were in the exponential growth phase at the start of exposure. Because infection alters filament mortality and population turnover, differences in population structure were minimised using the oligo-cell approach described below.

After confirming infection, the eight cultures were pooled separately by infection condition into two stocks (uninfected and infected) and distributed to 50 mL cell culture bottles for either MET-exposed or -unexposed treatments. Cultures were incubated as described in the previous section. MET stability in this experimental setup was previously confirmed, with negligible transformation observed [14]. The same experimental cultures were previously used to quantify chytrid fitness [14]

### Sample collection

From both MET-exposed and -unexposed cultures, three sample types were collected:

i) chytrid zoospores, ii) infected cyanobacterial filaments and iii) uninfected cyanobacterial filaments. The infected cyanobacterial filament samples contained both the cyanobacterial host and the infecting chytrid. Thus, the first two sample types represent the motile zoospore stage and the sessile infecting stage of *R. megarrhizum*, respectively. Zoospores and uninfected filaments were collected after 14 days of MET exposure, while infected filaments were collected after 15 days (21 days post-infection) due to the time required for manual collection.

To harvest zoospores, 1.5 mL of each infected culture was filtered through sterile 3 μm polycarbonate membranes to remove host filaments. To collect infected filaments, 4 mL of each infected culture were filtered through a sterile 10 μm nylon mesh. To collect uninfected filaments, 15 mL of each uninfected culture were filtered through a sterile 30 μm nylon mesh. Filtration removed long filaments, reducing entanglement and capillary clogging and, in infected samples, optimising the parasite-to-host ratio. Samples were collected using a micromanipulator (InjectMan; Eppendorf) equipped with a microinjector (CellTram Air; Eppendorf) and glass capillary attached to an inverted microscope (Nikon Ti Eclypse). Zoospores were collected at 400× magnification, and cyanobacterial filaments at 200× and 400×.

From each replicate, we collected 40–44 free-living chytrid zoospores, 6–11 infected cyanobacterial filaments (each containing approximately ten encysted zoospores or sporangia per filament side) and 30–34 uninfected cyanobacterial filaments. The number of filaments collected balanced two factors: minimising sampling time to reduce potential gene expression bias and obtaining sufficient material for oligo-cell RNA sequencing. The exact number of infections per filament could not be determined due to overlapping infections. One replicate per pollutant treatment and infection condition was sampled twice as a technical replicate, resulting in six samples per sample type. Samples were transferred to PCR tubes with 1–5 µL of medium, immediately shock-frozen in liquid nitrogen and stored at -80 °C.

### RNA sequencing

To assess the impact of MET on chytrid and cyanobacterial transcriptomes, total RNA was extracted and amplified at Qiagen (Hilden, Germany) using the QIAseq Single Cell RNA Library Kit UDI following the manufacturer’s instructions. Bacterial and fungal rRNA were depleted using the QIAseq FastSelect 5S/16S/23S kit (Qiagen) and the rRNA QIAseq FastSelect Yeast kit (Qiagen), respectively. The depletion was performed after cell lysis and gDNA removal and before reverse transcription, ligation and whole transcriptome amplification. For the infected filament samples containing both host and parasite, 1 µL of each FastSelect kit was used. For bacterial rRNA depletion, FastSelect was diluted 1:5 and the reverse-transcription primer volume doubled to 2 µL to minimise interference from oligonucleotide blockers. Due to the low starting material, additional purification steps were omitted.

Transcriptomes were sequenced at Macrogene Europe (Amsterdam, Netherlands) on the NovaSeq 6000 S4 platform (PE150, around 90 M reads/sample; Illumina). Read quality was scrutinised with FastQC v0.12.1 [27] and summarised via MultiQC v1.9 [28]. Reads were trimmed with Trimmomatic v0.39 [29] using LEADING: 3, TRAILING: 3, SLIDINGWINDOW: 4:15 and MINLEN: 36. Finally, rRNA sequences were removed using SortMeRNA v4.3.6 with the rRNA_databases_v4 [30]. Following rRNA removal, 36.18% of reads (*n* = 360 M) remained for chytrid zoospores, compared to 94.4% for uninfected cyanobacteria (*n* = 910 M) and 88% for infected filaments (*n* = 881 M).

### De novo transcriptome assembly and annotation for Rhizophydium megarrhizum Chy-Kol2008

To enrich the *R. megarrhizum* transcriptomic data obtained in this study, we combined our reads with those obtained from a bulk infected cyanobacterial culture [31]. Two *de novo* transcriptomes were assembled with Trinity [32]: one from zoospore samples only and one from infecting chytrid samples plus the bulk transcriptome. Non-fungal transcripts were removed using Diamond [33] BLASTX against the NCBI non-redundant protein (nr) database, and the retained fungal reads were reassembled using bedtools [34] and seqtk. Assembly quality was assessed with TrinityStats.pl and completeness with BUSCO (Benchmarking Universal Single-Copy Orthologs) [35] using the fungi_odb10 lineage dataset. Contamination was re- assessed using BLAST during annotation. Coding regions were predicted using TransDecoder [36] and annotated with Trinotate [37] using NCBI nr, UniProt, SwissProt [38], Pfam, Infernal [39], HMMER, signalP [40], TMHMM [41] and SQLite. The full workflow, tool versions and parameters are described in **Supplementary Methods** and **Figure S1.**

### Effect of MET and infection on gene expression

#### Transcript abundance estimation

To reduce incorrect mapping [42], a combined reference transcriptome was constructed, comprising the *de novo* generated *R. megarrhizum* transcriptome and predicted transcripts from the *P. agardhii* reference genome P. agardhii_No.976 (RefSeq assembly GCF_904830935.1; NCBI annotation release GCF_904830935.1-RS_2025_05_2; 98.02% completeness and 2.81% contamination). Transcript mapping and quantification were estimated using the Trinity align_and_estimate_abundance.pl script with Bowtie v1.3.1 and RSEM v1.3.3 abundance estimation method. Wrongly annotated transcripts (i.e. chytrid transcripts found in uninfected cyanobacteria or cyanobacterium transcripts found in zoospores) were removed from the count matrix. Separate transcript and gene count matrices were generated for chytrids and cyanobacteria.

#### Filtering low-quality transcripts and samples

Genes expressed in fewer than three samples per treatment were excluded from the analysis, along with repeated elements and known spurious-hit families. We calculated saturation levels, i.e. the number of genes detected with more than 0 counts with the sequencing depth of the sample, and with higher and lower simulated sequencing depths, with the R package NOISeq v2.52.0 [43] (**Figure S2**). Sequencing composition and gene detection were assessed for each sample (**Figure 2**). Based on the saturation plots and the chytrid-to- cyanobacterium read-count ratio, one sample was removed from the chytrid gene count matrix (met_chy_Z9), and four from the cyanobacterial matrix (control_both_In1, control_both_In4, control_both_In6, met_both_In10).

**Figure 2.**
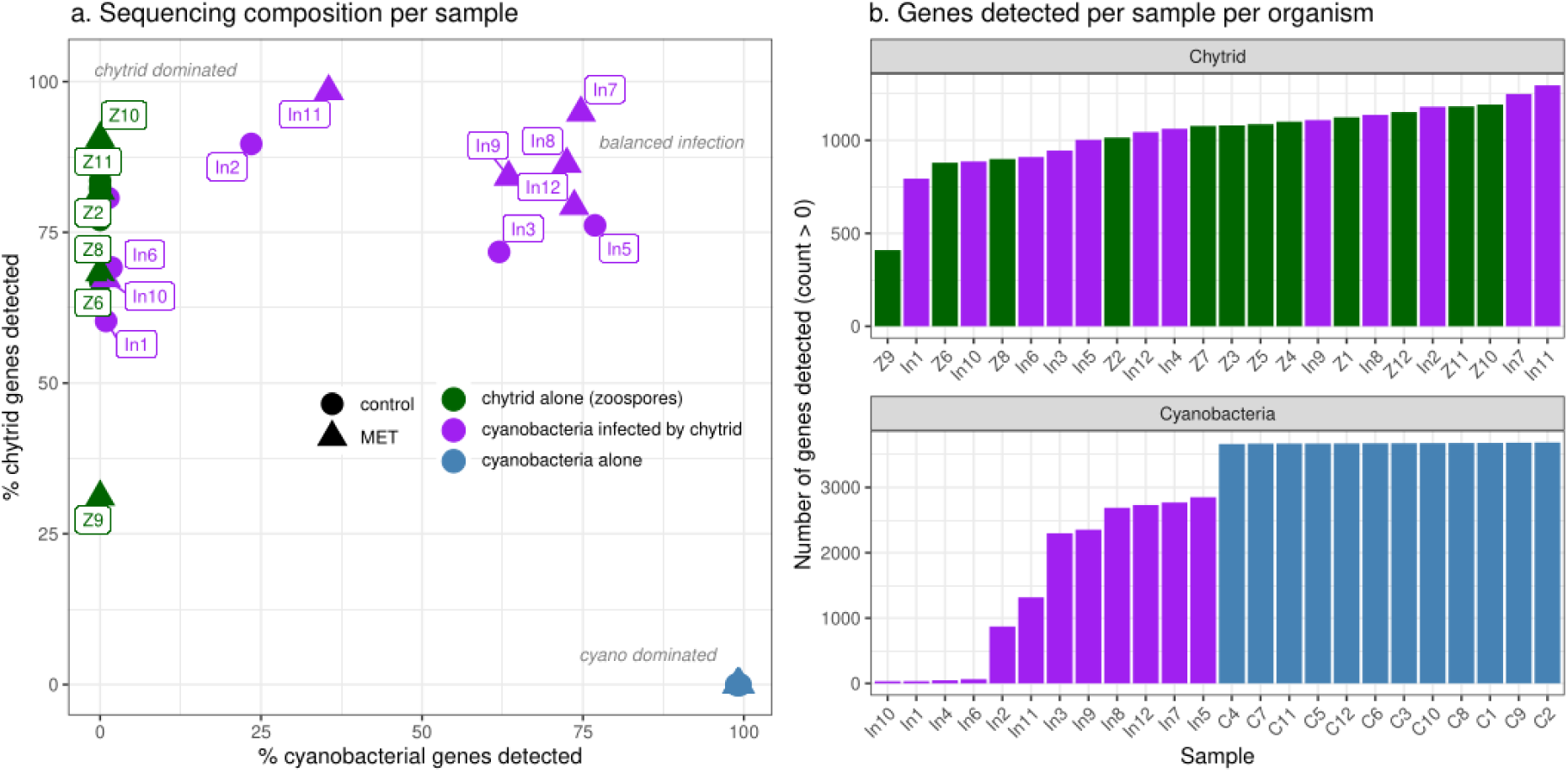
Sequencing composition and per-organism gene recovery across libraries. (a) Each point represents one library, positioned according to the percentage of cyanobacterial genes (x-axis) and chytrid genes (y-axis) detected (count > 0) among the total genes retained for each organism. Chytrid zoospore libraries recover chytrid but no cyanobacterial genes (top- left, "chytrid dominated") and uninfected cyanobacterial libraries the opposite (bottom-right, "cyano dominated"), whereas libraries of infected samples span a wide range of compositions, with a subset approaching balanced recovery of both transcriptomes ("balanced infection"). (b) Number of genes detected (count > 0) per library, separated by organism; libraries are ordered within each panel by the number of genes detected. Chytrid gene recovery overlaps broadly between zoospore and libraries containing both organisms, whereas cyanobacterial gene recovery collapses upon infection: libraries containing both organisms detect far fewer, and highly variable numbers of cyanobacterial genes than uninfected cyanobacterial libraries, which recover essentially the complete set (∼3,700 genes).

#### Differential gene expression analysis

Differential gene expression was analysed separately for chytrid and cyanobacterial transcriptomes using DESeq2 v1.48.1 [44]. Due to extreme differences in sequencing depth between single- and dual-species samples (size factors ranging from 360–440-fold when samples were pooled), contrasts were computed using condition-matched sample subsets rather than jointly across all conditions. Log2 fold changes were stabilised using ashr shrinkage [45] to account for variability in lowly expressed genes. Genes with a *padj* (FDR) < 0.05 were considered differentially expressed. Four contrasts were computed for chytrids: infection effects in the absence and presence of MET and MET effects on zoospores and infecting chytrids. For cyanobacteria, two contrasts tested MET effects on uninfected and infected filaments (**Table 1**); infection effects could not be reliably estimated due to low-quality filtering (**Figure 2**). For each pairwise comparison, genes with insufficient counts were removed using filterByExpr with default edgeR parameters [46]. Consequently, the number of genes included in each comparison was lower than the total number of detected genes.

**Table 1.** Overview of pairwise differential expression comparisons performed for chytrids and cyanobacteria.

| Conditions compared | Effect | N samples | N genes |
| --- | --- | --- | --- |
| **Chytrids** |  |  |  |
| Zoospores vs. infecting chytrids (No MET) | Life stage differences without MET | 12 | 882 |
| Zoospores vs. infecting chytrids (MET) | Life stage differences with MET | 11 | 1050 |
| MET-unexposed vs. MET-exposed zoospores | MET effect in free-living zoospores | 11 | 1012 |
| MET-unexposed vs. MET-exposed infecting chytrids | MET effect in infecting chytrids | 12 | 930 |
| **Cyanobacteria** |  |  |  |
| MET-unexposed vs. MET-exposed uninfected filaments | MET effect in uninfected cyanobacteria | 12 | 3512 |
| MET-unexposed vs. MET-exposed infected filaments | MET effect in infected cyanobacteria | 8 | 1943 |

#### Biomass loss versus infection effect on cyanobacteria

To test whether reduced gene detection in infected cyanobacteria libraries reflected lower biomass rather than infection-associated underexpression, we modelled expected detection under a biomass-only null model and evaluated gene frequency using a Poisson-binomial lower-tail test with Benjamini–Hochberg correction (for details, see **Supplementary Methods**; **Figure S3**).

#### Cyanobacterium and chytrid gene co-expression analyses

Weighted gene co-expression networks were constructed separately for each organism (WGCNA) [47], using only single-species samples to avoid composition-driven cross-species correlations: 12 uninfected-cyanobacterial libraries (3,706 genes) and 11 chytrid zoospore libraries (1,252 genes after quality filtering). Signed networks were built from variance-stabilised counts using biweight midcorrelation (maxPOutliers = 0.1), with the lowest soft-thresholding power achieving a scale-free fit of R² ≥ 0.85. Module eigengenes were detected by dynamic tree cutting (minModuleSize = 30, deepSplit = 2, mergeCutHeight = 0.25) and correlated with MET using Pearson correlation with Benjamini–Hochberg correction. Differentially expressed genes (DEGs) were mapped to modules, and module enrichment was tested using one-sided Fisher’s exact tests (BH-corrected).

Plots were generated using the R packages ggplot2 v3.5.1 [48], igraph v2.1.4 [49], tidygraph v1.3.1 and ggraph v2.2.1 and the software Inkscape v1.2 [50]. All scripts are archived on Zenodo [51]. Calculations were made on the high-performance computer at Freie Universität Berlin [52].

## Results

### De novo transcriptome assembly of Rhizophydium megarrhizum Chy-Kol2008

The assembled transcriptome contained 16,909 transcripts and 3,894 Trinity “genes”, with a GC content of 40.14%. The N50 was 7,701 bp for all transcripts and 11,176 bp for the longest isoforms. Transcriptome completeness showed 68.3% complete BUSCOs out of 758 BUSCO groups searched, of which 9.9% were single-copy and 58.4% were duplicated. Fragmented BUSCOs accounted for 9.6%, while 22.0% of BUSCOs were missing. After sample quality filtering based on outlier detection and sequencing saturation (**Figure 2**), RNAseq data for 1,317 chytrid genes and 3,706 cyanobacteria genes were retained for downstream analyses (**Table 1**).

### Dual-RNAseq of the host–parasite system

#### Chytrid gene expression across life cycle stages and under MET exposure

Across all four comparisons, infection and MET exposure affected a limited number of high-confidence genes associated with distinct processes (**Figure 3**, **Table 2**), suggesting specific physiological adjustments rather than a coordinated response. In the absence of MET, seven genes were differentially expressed between infecting chytrids and zoospores (control_chy vs. control_both; FDR < 0.05; **Figure 3a**, e, **Table S2**). Genes encoding an ABC transporter (ABCB6; *padj* = 0.0075, log2FC = -2.6) and a cytosolic carboxypeptidase 1 (CBPC1; *padj* = 0.029, log2FC = -2.1) were underexpressed, whereas those encoding α-amylase (AMYA; *padj* = 0.012, log2FC = 2.5), Rho GDP-dissociation inhibitor (GDIR; *padj* = 0.0075, log2FC = 3.3), replication factor (RFC4; *padj* = 0.029, log2FC = 2.4), tRNA-splicing ligase (TRNL; *padj* = 0.029, log2FC = 1.5) and N-acetyltransferase (ACA1; *padj* = 0.031, log2FC = 2.1) were overexpressed.

**Figure 3.**
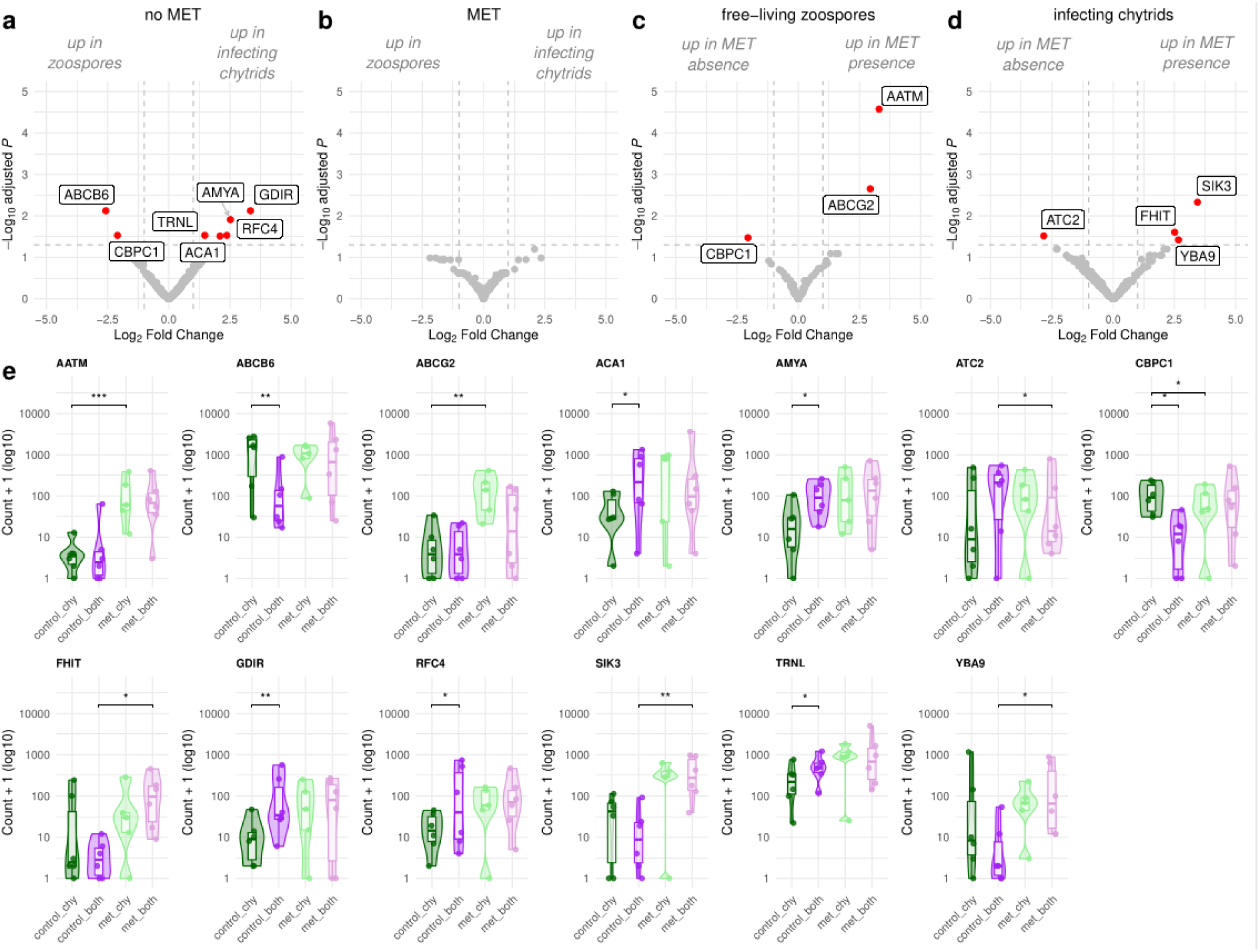
Interactive effects of metolachlor (MET) and infection on chytrid gene expression. (a–d) Volcano plots of the four pairwise DESeq2 contrasts, showing log₂ fold change against -log₁₀ adjusted *p*-value. Red points indicate significantly differentially expressed genes (adjusted *p* < 0.05), with underexpressed genes to the left and overexpressed genes to the right; grey points indicate genes that are not significantly differentially expressed. Dashed lines indicate the |log₂FC| = 1 and adjusted *p* = 0.05 thresholds. The baseline condition is shown on the left and the comparison condition on the right: positive log₂ fold changes (right) indicate overexpressed genes in the comparison condition relative to baseline, and negative log₂ fold changes (left) underexpressed genes in the comparison condition relative to baseline. **(e)** Raw expression (count + 1, log₁₀ scale) of every differentially expressed gene across the four experimental conditions: zoospores without (control_chy) and with MET (met_chy), and infecting chytrids without (control_both) and with MET (met_both). Violins show the distribution, boxplots the median and interquartile range, and points the individual libraries; brackets and asterisks indicate the contrast(s) in which the gene was significant (* adjusted p < 0.05, ** < 0.01, *** < 0.001).

**Table 2.**
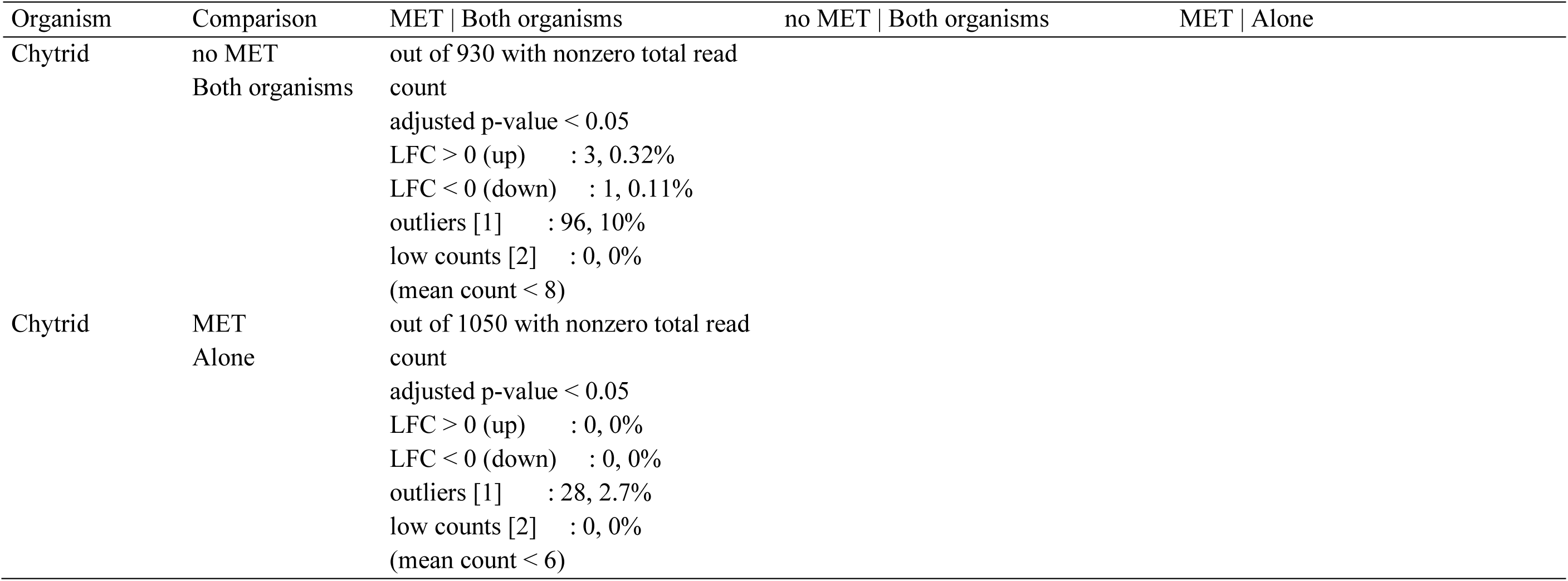
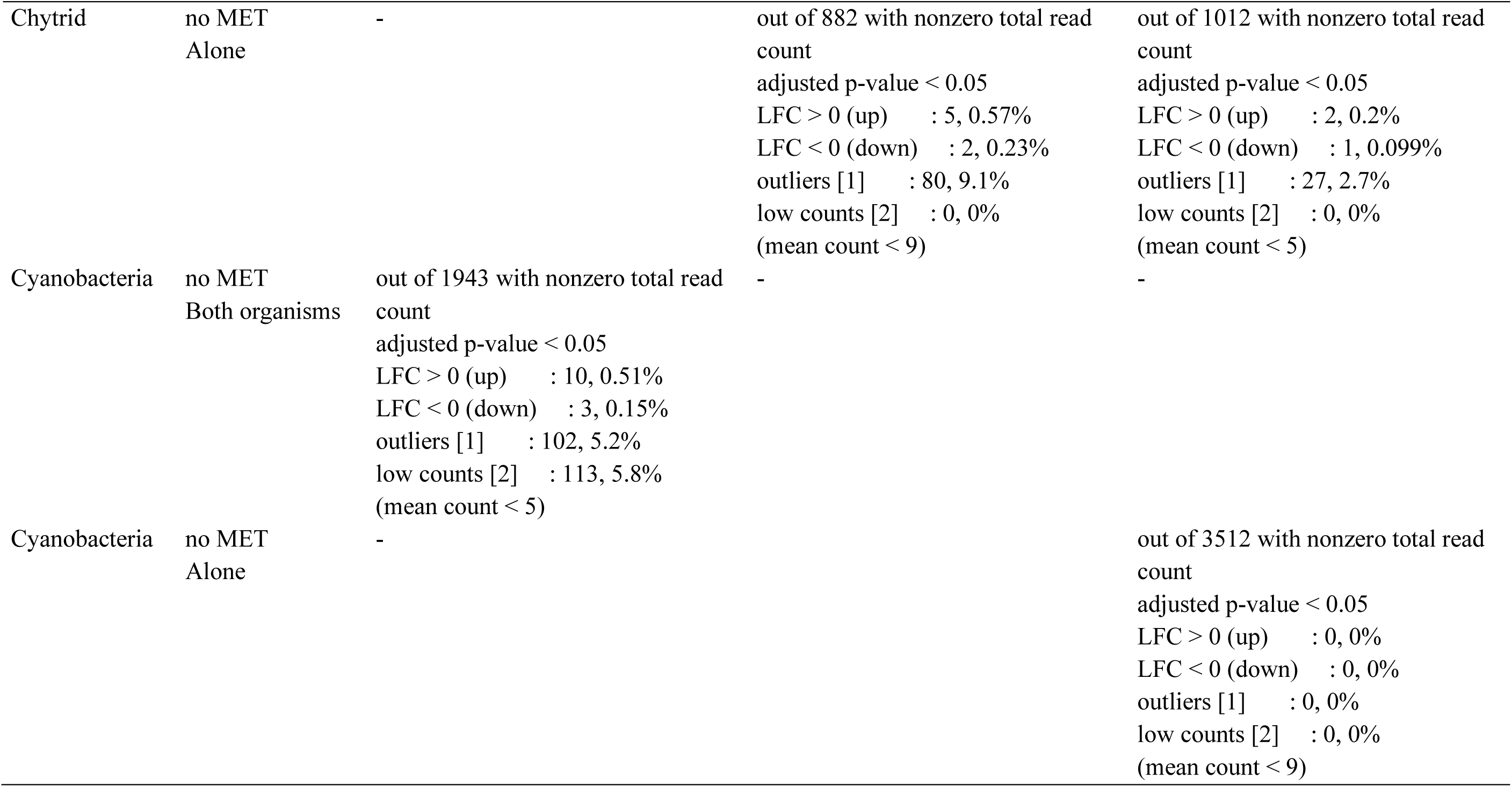
Summary of differentially expressed genes (DEGs) in chytrids and cyanobacteria under different experimental conditions. Pairwise contrasts were performed to assess the effect of MET, infection status (chytrids or cyanobacteria alone vs. infection) and their interaction. For each comparison, the number and percentage of transcripts with significant differential expression are reported (absolute log2 fold change ≥ 1, adjusted *p* < 0.05, number of outliers, low count genes, and mean count, i.e. the optimal mean count threshold for removing genes that have very low power to be detected as significant).

MET exposure altered gene expression in both zoospores and infecting chytrids compared to their unexposed counterparts. In zoospores, one gene encoding a cytosolic carboxypeptidase 1 (CBPC1; *padj* = 0.034, log2FC = -2.1) was underexpressed, whereas genes encoding aspartate aminotransferase (AATM; *padj* = 0.000026, log2FC = 3.3) and ABC transporter G family member 2 (ABCG2; *padj* = 0.0022, log2FC = 2.9) were overexpressed (control_chy vs. met_chy; FDR < 0.05; **Figure 3c**, e, **Table S2**). The gene encoding CBPC1 was the only gene consistently underexpressed during infection in the absence of MET and in MET-exposed zoospores (**Figure 3a, c**). In infecting chytrids, the gene encoding calcium-transporting ATPase 2 (ATC2; *padj* = 0.03, log2FC = -2.8) was underexpressed, whereas genes encoding a serine/threonine-protein kinase (SIK3; *padj* = 0.0047, log2FC = 3.4), a homologue of the fragile histidine triad protein (FHIT; *padj* = 0.025, log2FC = 2.5) and an uncharacterised protein (YBA9; *padj* = 0.038, log2FC = 2.7; **Figure 3d**, e, **Table S2**) were overexpressed.

No genes were differentially expressed between infecting chytrids and zoospores in the presence of MET (met_chy vs. met_both: no significant genes; **Figure 3b**, **e**). However, the expression pattern of the AATM- and ABCG2-encoding genes, which were significantly overexpressed in MET-exposed zoospores relative to unexposed ones, was similar in MET-exposed infecting chytrids vs MET-exposed zoospores, but not significant. Similarly, SIK3-, FHIT- and YBA9-encoding genes were significantly overexpressed in MET-exposed infecting chytrids compared to their unexposed counterparts and showed the same trend in MET-exposed infecting chytrids relative to MET-exposed zoospores (**Figure 3e**).

#### Cyanobacterial gene expression in response to MET and chytrid infection

In uninfected cyanobacteria, MET produced no detectable transcriptional response, with no genes passing the significance threshold and all fold-changes clustering around zero (**Figure 4a**, **Table 2**). In contrast, 13 genes were differentially expressed in MET-exposed infected cyanobacteria compared with their unexposed counterparts (control_both vs. met_both, FDR < 0.05; **Figure 4b**, **Table S2**). However, as most genes showed very low read counts close to the detection limit (**Figure 4c**), these results should be interpreted with caution, as some changes may reflect limited transcript coverage rather than biological responses.

**Figure 4.**
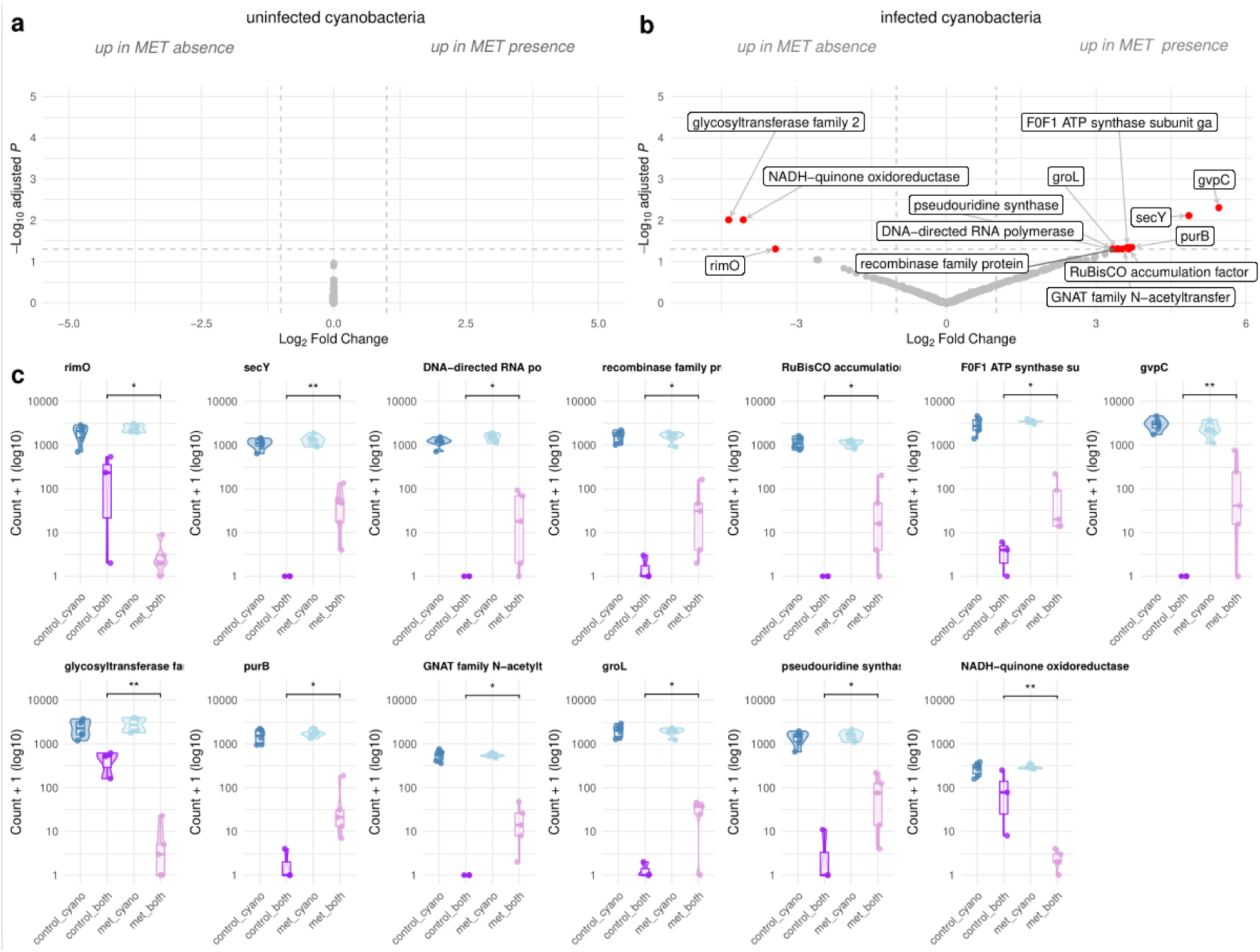
Effects of metolachlor (MET) on cyanobacteria gene expression. (a, b) Volcano plots of the two pairwise DESeq2 contrasts, showing log₂ fold change against -log₁₀ adjusted *p*-value. Red points indicate significantly differentially expressed genes (adjusted *p* < 0.05), with underexpressed genes to the left and overexpressed genes to the right; grey points indicate genes that are not significantly differentially expressed. Dashed lines indicate the mark |log₂FC| = 1 and adjusted *p* = 0.05 thresholds. The baseline condition is shown on the left and the comparison condition on the right: positive log₂ fold changes (right) indicate overexpressed genes in the comparison condition relative to baseline, and negative log₂ fold changes (left) underexpressed genes in the comparison condition relative to baseline. **(c)** Raw expression (count + 1, log₁₀ scale) of every differentially expressed gene across the four experimental conditions: uninfected cyanobacteria without (control_cyano) and with MET (met_cyano), and infected cyanobacteria without (control_both) and with MET (met_both). Violins show the distribution, boxplots the median and interquartile range, and points the individual libraries; brackets and asterisks indicate the contrast(s) in which the gene was significant (* adjusted p < 0.05, ** < 0.01, *** < 0.001).

Four DEGs showed consistently detectable reads irrespective of MET exposure: genes encoding 30S ribosomal protein S12 methylthiotransferase (*rimO*, 77286364; *padj* = 0.049, log2FC = -3.4), glycosyltransferase (77287779; *padj* = 0.0098, log2FC = -4.4) and NADH-quinone oxidoreductase subunit J (77289563, *ndhJ*; *padj* = 0.0098, log2FC = -4.1) were underexpressed, whereas the gene encoding the F0F1 ATP synthase γ-subunit was overexpressed (77287231; *padj* = 0.045, log2FC = 3.6). The remaining nine DEGs included those encoding the preprotein translocase subunit SecY (77286711; *padj* = 0.0077, log2FC = 4.9), DNA-directed RNA polymerase subunit α (77286717; *padj* = 0.049, log2FC = 3.3) and a gas-vesicle protein (*gvpC*; 77287375; *padj* = 0.005, log2FC = 5.5).

#### Biomass loss versus infection effects on cyanobacterial gene expression

Of the 3,536 genes reliably expressed in uninfected cyanobacteria, only one gene, encoding a glutathione S-transferase, met the criteria for an infection-specific loss rather than loss due to reduced cyanobacterial biomass and the lower proportion of cyanobacterial reads in the dual-transcriptome libraries (**Figure S3**). This gene was detected at low abundance only in the deepest sequenced dual-transcriptome libraries, whereas the remaining candidate genes were more highly abundant and therefore more sensitive to sequencing depth. A complementary compositional differential expression analysis was likewise uninformative because the DESeq2 size factors were strongly confounded with infection status (alone ≈ 4–7.5, co-culture ≈ 0.01– 0.26), violating the assumption that most genes remain unchanged and making estimates unreliable in the read-limited dual-transcriptome libraries. Together, these analyses indicate that the low and variable cyanobacterial fraction recovered in infected-cyanobacteria samples prevented reliable distinction between infection-associated regulation and changes resulting from reduced biomass.

#### Gene co-expression analyses of cyanobacteria and chytrids

The cyanobacterial network reached the scale-free topology criterion (power = 15, R² = 0.85) and resolved 64 modules (**Figure 5a**), but no module eigengene was associated with MET (minimum *padj* = 0.33). The 13 DEGs were distributed across 11 modules (**Figure 5b**). In contrast, the chytrid network showed weak structure, with a maximum scale-free topology fit of R² = 0.69 (**Figure 5c**). The 10 DEGs were distributed across six modules, with two genes unassigned (**Figure 5d**). As the chytrid network did not meet the scale-free criterion, these assignments are exploratory.

**Figure 5.**
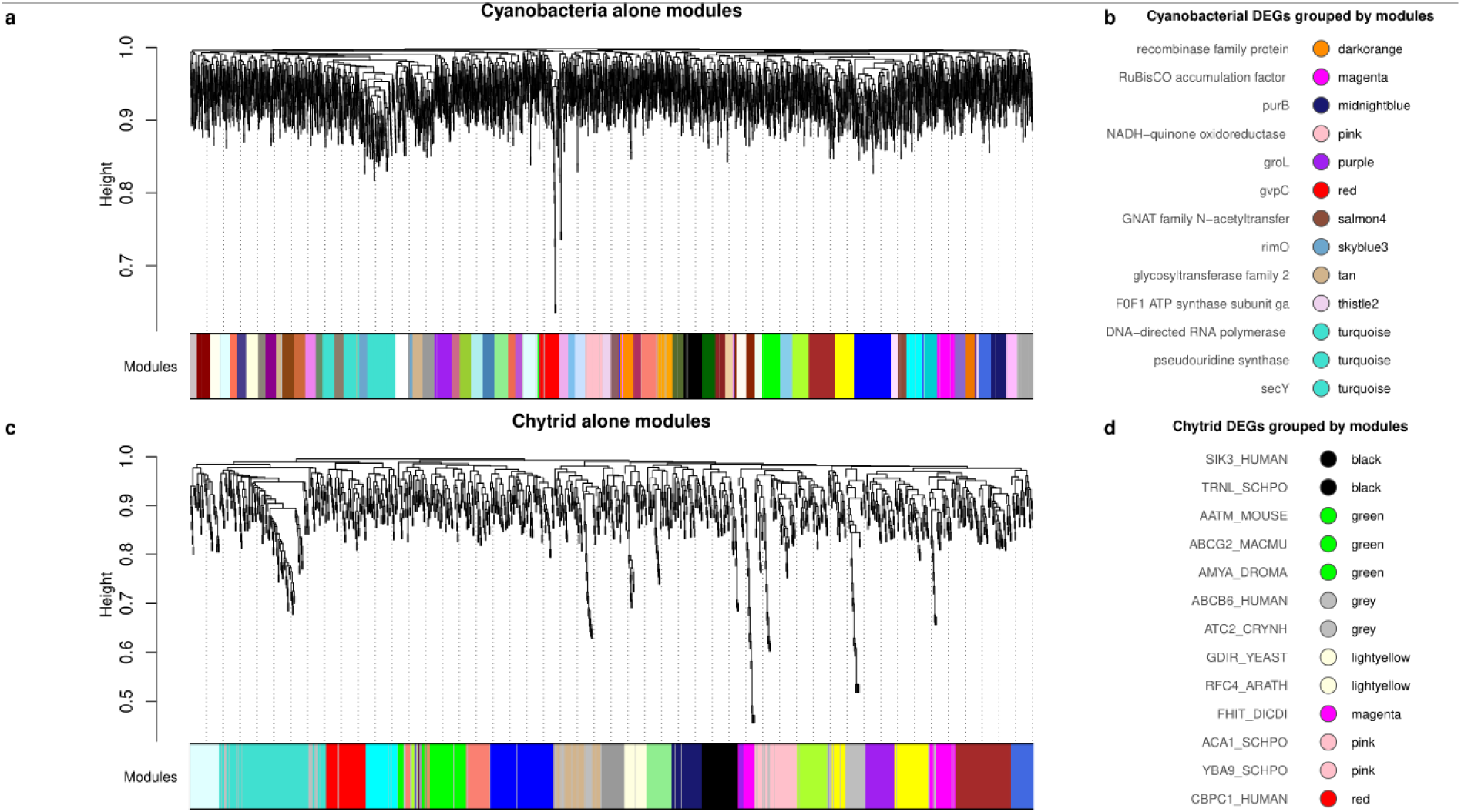
Co-expression networks for each organism and distribution of differentially expressed genes across modules. Signed weighted gene co-expression networks (WGCNA, biweight midcorrelation) were constructed separately for each organism, using single-organism libraries to avoid composition-driven correlations in multi-organism samples. Variance- stabilised counts were used as input. (a) Hierarchical clustering dendrogram of the 3,706 genes in the cyanobacterial network (12 libraries); the colour bar beneath shows the dynamic-tree-cut module assignment (64 modules; soft-thresholding power 15, scale-free fit R² = 0.85). (b) Module membership of the 13 cyanobacterial DEGs (the metolachlor effect in infected cyanobacteria), each shown as a point coloured by its WGCNA module, with the module name at the right and the protein product at the left. WGCNA identifies modules as clusters of genes with correlated expression patterns that are likely involved in shared functions. The DEGs are spread across 11 modules; only the turquoise module contains more than one (the SecY translocase, the DNA-directed RNA polymerase α subunit and a pseudouridine synthase). No module was significantly enriched for DEGs, and no module eigengene was correlated with metolachlor, after FDR correction. (c) As in (a) for the chytrid network (1,252 genes, 11 libraries). This network did not reach the scale-free topology criterion (maximum R² = 0.69; default power 12) and was only weakly modular. (d) Module membership of the 13 chytrid DEGs, coloured by WGCNA module (UniProt entry names; grey = genes not assigned to any module). Two genes were unassigned; the green (AATM, ABCG2, AMYA), lightyellow (RFC4, GDIR), black (SIK3, TRNL) and pink (ACA1, YBA9) modules each contain two or more DEGs, but no module was significantly enriched after FDR correction. .

Despite the absence of significant module-level associations with MET, several DEGs co-occurred within modules. In cyanobacteria, only one module contained multiple DEGs, encoding SecY translocase, DNA-directed RNA polymerase α-subunit and pseudouridine synthase; the remaining DEGs occurred in separate modules. In chytrids, genes encoding AATM, ABCG2 and AMYA co-occurred in one module, while genes encoding the infection-associated RFC4 and GDIR, SIK3 and TRNL, and ACA1 and YBA9 each co-occurred within individual modules.

## Discussion

To our knowledge, this study is the first to provide a replicated, experimental, composition-aware dual-transcriptome analysis designed to resolve how pollutant exposure affects the interaction between *P. agardhii* and *R. megarrhizum*. MET primarily affected the chytrid parasite, altering host**–**parasite dynamics while exerting little direct transcriptional effect on the cyanobacterial host. Our results demonstrate that herbicide effects depend strongly on the ecological context in which organisms interact.

### Physiological transition from zoospores to infecting stage

The transcriptional changes observed in infecting chytrids indicate a coordinated shift from a motile dispersal stage towards an actively growing parasitic stage. Reduced expression of the gene encoding cytosolic carboxypeptidase 1 may reflect this transition, given its role in tubulin modification, microtubule organisation and ciliary or flagellar function [53]. Conversely, increased expression of the gene encoding Rho GDP-dissociation inhibitor may indicate alterations in cytoskeletal reorganisation, membrane trafficking and polarised growth [54]. Rho GDP-dissociation inhibitors are also involved in regulating the yeast-to-hyphae switch in *Candida* [55]. Given that rhizoid morphogenesis in chytrids resembles hyphal growth [56], these changes may be related to rhizoid development and host colonisation, consistent with cytoskeletal genes being highly expressed during the host-associated stage of *R. megarrhizum* [57]. The overexpression of genes encoding N-acetyltransferase and α-amylase may further reflect the metabolic demands of the infecting stage, as these enzymes have been implicated in fungal infection and colonisation [58–60]. Similarly, increased expression of genes encoding RFC4 and TRNL may reflect higher demands for DNA synthesis and RNA processing during parasite development [61–63].

### MET exposure disrupts the developmental transition of chytrids

The transcriptional responses indicate that MET affects both chytrid life stages. In free-living zoospores, MET induced transcriptional changes that suggest interference with the normal developmental programme. Reduced expression of the gene encoding cytosolic carboxypeptidase 1 may indicate altered cytoskeletal remodelling associated with the flagellated stage. As this gene was also underexpressed in infecting chytrids compared with zoospores in the absence of MET, its reduced expression under MET exposure suggests a shift towards a transcriptional state associated with the sessile stage prior to host contact. Additionally, overexpression of genes encoding AATM, involved in amino acid and carbon metabolism, and ABCG2, involved in xenobiotic transport [64], may reflect metabolic adjustment in response to MET exposure. The co-occurrence of infection- and MET-responsive genes within network modules is consistent with an altered transcriptional programme during the zoospore-to-sporangium transition.

In contrast, MET predominantly altered genes involved in regulatory processes in infecting chytrids. Underexpression of the gene encoding ATC2 suggests altered calcium homeostasis during MET exposure, which may affect parasite development given the role of calcium signalling in fungal adaptation and chytrid reproduction [65, 66]. MET also increased the expression of genes encoding SIK3, involved in cell proliferation, differentiation and signalling in fungi [67], as well as FHIT and the uncharacterised protein YBA9. Although their functions in chytrids remain uncertain, FHIT has been linked to cell proliferation and survival [68], while YBA9 belongs to the ORM/ORMDL family of membrane-bound regulators associated with sphingolipid biosynthesis. Together these responses suggest altered cellular and lipid regulation during MET exposure.

Although only a limited number of genes were significantly differentially expressed in response to MET in zoospores, several genes overexpressed during infection in the absence of MET (e.g. AMYA, GDIR, RFC4 and TRNL) displayed the same directional trend in MET-exposed zoospores. This pattern suggests that MET partially shifts the transcriptional state of zoospores towards that of infecting chytrids, thereby reducing the molecular distinction between the dispersal and infection stages. This convergence may reshape host–parasite interactions and reduce chytrid virulence, consistent with our previous observations that long-term MET exposure reduced chytrid fitness and delayed epidemic onset in cyanobacterial cultures [14]. These findings support our first hypothesis that MET alters the expression of genes associated with host colonisation and nutrient uptake in zoospores.

### MET alters cyanobacterial gene expression only during exposure to chytrid infection

In contrast to the pronounced transcriptional responses observed in the chytrid, MET did not induce significant transcriptional changes in uninfected cyanobacteria. At the network level, MET-responsive genes were scattered across modules, with no evidence of a coordinated network-level response. This differs from the growth-promoting effect observed during short-term MET exposure [14]. However, this growth promotion was observed after seven days of exposure, whereas the present samples were collected after 14 days. Thus, transcriptional responses associated with the growth-promoting effect may no longer have been detectable at the later sampling time.

Among the differentially expressed genes detected in MET-exposed infected filaments, reduced expression of a glycosyltransferase-encoding gene involved in polysaccharide biosynthesis [69] and of *ndhJ*, involved in respiration and photosynthesis [70], suggests altered carbohydrate and energy metabolism. Increased expression of the gene encoding the F0F1 ATP synthase γ-subunit may represent a compensatory response, while reduced *rimO* expression may indicate altered ribosome maturation and protein synthesis [71]. Although these results are tentative given the limited cyanobacterial transcript recovery, they suggest subtle metabolic adjustments under combined MET exposure and chytrid infection.

MET exposure was also associated with increased expression of *gvpC*, encoding a gas-vesicle protein involved in cyanobacterial buoyancy [72] and regulation of the surface area-to-volume ratio [73, 74]. This change might reflect adjustments in cellular positioning and interaction with the surrounding environment. Bulk transcriptomic analyses of *P. agardhii* during *R. megarrhizum* infection also identified changes in gas-vesicle-associated genes [57]. Although the datasets cannot be directly compared, the recurrence of this response suggests that buoyancy-related processes may be associated with the physiological response to chytrid infection.

Altogether, our findings are consistent with previous physiological observations. Using a multi-biomarker approach, we found that MET alone did not induce detectable physiological stress in *P. agardhii*, whereas simultaneous MET exposure and chytrid infection caused a moderate stress response [24]. Despite the different exposure period, both approaches showed that MET alone had little detectable impact on the cyanobacterial host, whereas molecular and physiological responses emerged under co-exposure to MET and parasites. This consistency supports the apparent resilience of *P. agardhii* to MET in the absence of its parasite and suggests that chytrid infection can modulate the cyanobacterial response to pollutant exposure. These results partially support our second hypothesis: although no oxidative-stress-related genes were overexpressed, transcriptional changes were detected only in MET-exposed infected cyanobacteria.

### Study limitations

The small number of DEGs detected in the present study likely reflects a combination of biological, technical and analytical factors. Biologically, chytrid infection strongly affects cyanobacterial biomass and may degrade host genetic material during parasite development. Infection is also heterogeneous within cyanobacterial filaments, with infected and uninfected cells co-existing within the same sample, potentially diluting subtle transcriptional signals.

Moreover, strong infection-associated responses may mask additional effects of MET in infected cyanobacteria.

Technical factors may further influence gene detection, including differences in biomass abundance, genome complexity and genome size between cyanobacteria and chytrids, incomplete cell lysis (particularly of chytrids) and limited removal of chytrid rRNA (decreasing the proportion of mRNA reads) due to the low specificity of the yeast rRNA removal kit. Analytical factors, including the stringent thresholds applied for gene selection and differential expression analysis, may have further reduced the number of detected genes. Stringency was necessary to ensure robust biological signals, but the absence of differential expression does not necessarily indicate the absence of biological effects, as responses can fall below the detection threshold or be masked by infection-associated processes.

### Ecological implications

Our results demonstrate that herbicide pollution can reshape host–parasite interactions, with consequences beyond direct toxicity. By weakening chytrid-mediated top-down control, MET may indirectly favour cyanobacterial bloom persistence. As chytrids transfer energy from inedible phytoplankton to higher trophic levels via the mycoloop, pollutant-induced changes in host–parasite interactions may cascade through aquatic trophic webs, affecting energy flow and ecosystem functioning. These findings highlight the importance of considering pollutant effects on microbial interactions, rather than only on individual species. Future studies should determine how such pollutant-induced changes translate into community- and ecosystem-level responses.

## Funding sources

This work was supported by the Deutsche Forschungsgemeinschaft (DFG) grants MA 9934/1- 1, project no. 471387895 (E.B.M.R.) and STR1349/2-1/2-2, project no. 432453260 (J.F.H.S.), and by the European Commission HORIZON–MSCA-2022-DN program grant 101120280 (J.W.). Computing resources for the bioinformatic analyses were partially funded by the German Federal Ministry of Education and Research (BMBF, Förderkennzeichen 033W034A).

## Data availability

All the raw data generated in this project, along with the de novo transcriptome assembly, have been deposited in the European Nucleotide Archive (ENA) database under the project accession number PRJEB98683.

## Conflict of interest disclosure

The authors declare no conflicts of interest.

## Authors’ contribution

**EBMR**: Conceptualisation, Investigation (lead), Formal analysis, Funding acquisition, Methodology (lead of experimental work), Resources, Project administration, Visualisation, Writing - Original Draft, Writing - Review & Editing; **AB**: Data curation, Formal analysis (lead of bioinformatics), Methodology (lead of bioinformatics), Software, Visualisation (lead), Writing - Review & Editing; **JFHS**: Formal analysis (supporting), Funding acquisition, Investigation (supporting), Methodology (lead of molecular work), Resources, Writing - Review & Editing; **JW**: Resources, Writing - Review & Editing.

## Supplementary Methods

### De novo transcriptome assembly and annotation for Rhizophydium megarrhizum Transcriptome assembly

To enrich the *R. megarrhizum* transcriptomic data obtained in this study, we incorporated RNA sequences from a bulk infected cyanobacterial culture containing both chytrid zoospores and infecting chytrids, as well as their cyanobacterial hosts [1]. This culture was incubated under the conditions described in the subsection “*Cyanobacteria and chytrids culturing conditions”*, with cell collection and RNA sequencing described in [2]. The full bioinformatics workflow is shown in **Figure S1**. Reads from the 25 chytrid-containing samples were concatenated into two subsets: 12 zoospore samples (188 M paired-end reads) and 12 oligo-cell samples of chytrids infecting cyanobacteria plus one bulk culture sample (487.3 M paired-end reads). Two *de novo* transcriptomes were assembled using Trinity v2.15.2 [3] with default parameters and -- normalize_by_read_set.

Contaminants (transcripts unlikely to belong to *R. megarrhizum*) were filtered out using Diamond v2.1.8 [4] BLASTX search against the NCBI non-redundant protein database (--sensitive --max-target-seqs 1 --evalue 1e-5). Transcripts mapping to the Fungi kingdom were retained (*n* = 43,644, 7.69% for the zoospore assembly; *n* = 36,112, 4.35% for the chytrids infecting cyanobacteria assembly). Reads aligning to the original transcriptomes with minimap2 v2.29 [5] were filtered for paired-end reads mapping to the fungal transcripts using bedtools v2.31.0 [6] and seqtk v1.4. The resulting 48.9 M paired-end reads were assembled with Trinity as previously described. Contamination was re-assessed using BLAST during annotation with Trinotate (see below). The final *R. megarrhizum* chytrid assembly retained transcripts with best hits to eukaryotes (50.3% of the total transcripts), while cyanobacterial transcripts were removed. Transcriptome quality was assessed with TrinityStats.pl and completeness was assesses with BUSCO (Benchmarking Universal Single-Copy Orthologs) v5.8.2 [7] using the fungi_odb10 lineage dataset.

### Transcriptome annotation

Coding regions of the decontaminated transcriptome were predicted using TransDecoder v5.7.1 and annotated with Trinotate v4.0.2 [8]. TransDecoder peptides were searched against the NCBI nr database using Diamond BLASTP (--sensitive --max-target-seqs 10 --max-hsps 1 --evalue 1e-5). Trinotate was run with --run “swissprot_blastp swissprot_blastx pfam infernal” and --use-diamond for homology searches against UniProt Swiss-Prot database [9]. The “pfam” option was used for protein domain identification with profile hidden Markov models using HMMER v3.4, while the “infernal” option was applied for noncoding RNA identification with Infernal v1.1.5 [10]. Signal peptides and their cleavage sites, as well as transmembrane helices were predicted using signalP v6.0 [11] and TMHMM v2.0 [12], respectively. All information was stored in an SQLite v3.43.1 database later used for annotation by Trinotate.

### Biomass loss versus infection effect on cyanobacteria

We modelled gene detection in infected-cyanobacteria libraries under a biomass-only null model to determine whether reduced detection relative to uninfected libraries could be explained by lower cyanobacterial biomass and sequencing depth or by infection-associated underexpression. For each gene reliably expressed in uninfected cyanobacteria (≥ 5 reads in ≥ 80% of uninfected libraries; *n* = 3,536), we calculated its mean relative abundance and, together with the cyanobacterial sequencing depth of each infected-cyanobacteria library, estimated its expected read count and detection probability (≥5 reads; negative binomial; size = 1). We then tested whether genes were detected less frequently than expected using a Poisson-binomial lower-tail test with Benjamini–Hochberg correction. A gene was classified as an infection-linked loss only if all four criteria were met: (i) significantly lower detection frequency than expected (*padj* < 0.05); (ii) decrease in relative abundance (log₂ fold-change < −1); (iii) expected detection in ≥70% in infected-cyanobacteria libraries; and (iv) detection in no more than one infected library (**Figure S3**).

## Supplementary Tables

**Table S1.** Experimental design.

| Sample ID | Organism | Number of cells/filaments | Treatment | Culture |
| --- | --- | --- | --- | --- |
| Z1 | Zoospores (chytrid) | 40 | control | Infected |
| Z2 | Zoospores (chytrid) | 40 | control | Infected |
| Z3 | Zoospores (chytrid) | 41 | control | Infected |
| Z4 | Zoospores (chytrid) | 40 | control | Infected |
| Z5 | Zoospores (chytrid) | 42 | control | Infected |
| Z6 | Zoospores (chytrid) | 41 | control | Infected |
| Z8 | Zoospores (chytrid) | 43 | MET | Infected |
| Z9 | Zoospores (chytrid) | 42 | MET | Infected |
| Z10 | Zoospores (chytrid) | 42 | MET | Infected |
| Z11 | Zoospores (chytrid) | 43 | MET | Infected |
| Z7 | Zoospores (chytrid) | 43 | MET | Infected |
| Z12 | Zoospores (chytrid) | 44 | MET | Infected |
| In1 | Infected_filaments (both) | 6 | control | Infected |
| In2 | Infected_filaments (both) | 6 | control | Infected |
| In3 | Infected_filaments (both) | 7 | control | Infected |
| In4 | Infected_filaments (both) | 6 | control | Infected |
| In5 | Infected_filaments (both) | 6 | control | Infected |
| In6 | Infected_filaments (both) | 8 | control | Infected |
| In7 | Infected_filaments (both) | 6 | MET | Infected |
| In8 | Infected_filaments (both) | 8 | MET | Infected |
| In9 | Infected_filaments (both) | 9 | MET | Infected |
| In10 | Infected_filaments (both) | 11 | MET | Infected |
| In11 | Infected_filaments (both) | 7 | MET | Infected |
| In12 | Infected_filaments (both) | 8 | MET | Infected |
| C1 | Filaments (cyanobacteria) | 30 | control | Uninfected |
| C3 | Filaments (cyanobacteria) | 34 | control | Uninfected |
| C4 | Filaments (cyanobacteria) | 34 | control | Uninfected |
| C5 | Filaments (cyanobacteria) | 30 | control | Uninfected |
| C2 | Filaments (cyanobacteria) | 32 | control | Uninfected |
| C6 | Filaments (cyanobacteria) | 32 | control | Uninfected |
| C7 | Filaments (cyanobacteria) | 30 | MET | Uninfected |
| C8 | Filaments (cyanobacteria) | 32 | MET | Uninfected |
| C9 | Filaments (cyanobacteria) | 30 | MET | Uninfected |
| C10 | Filaments (cyanobacteria) | 31 | MET | Uninfected |
| C11 | Filaments (cyanobacteria) | 30 | MET | Uninfected |
| C12 | Filaments (cyanobacteria) | 30 | MET | Uninfected |

Identical colour = replicates from the same culture

**Table S2.**
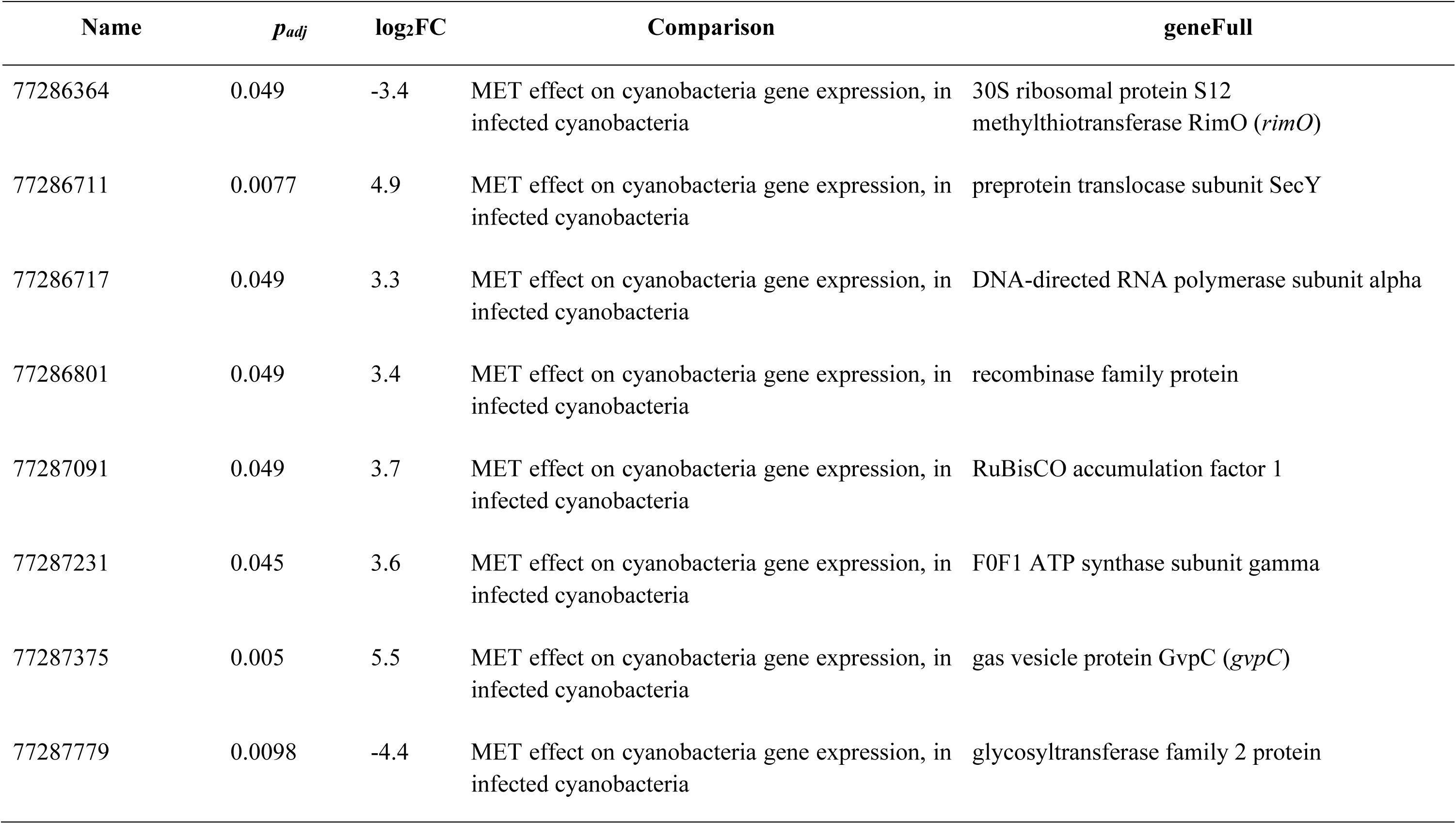
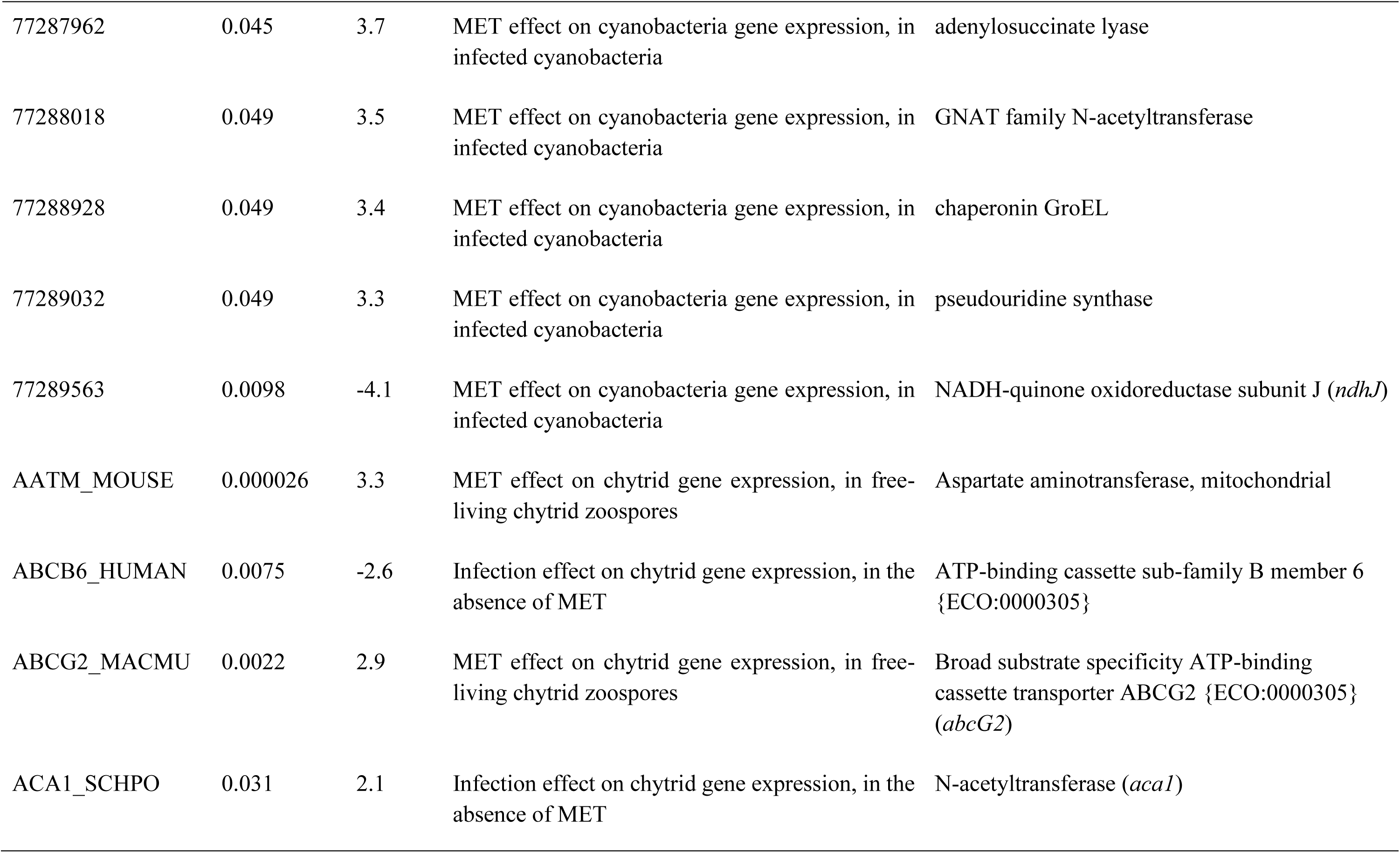
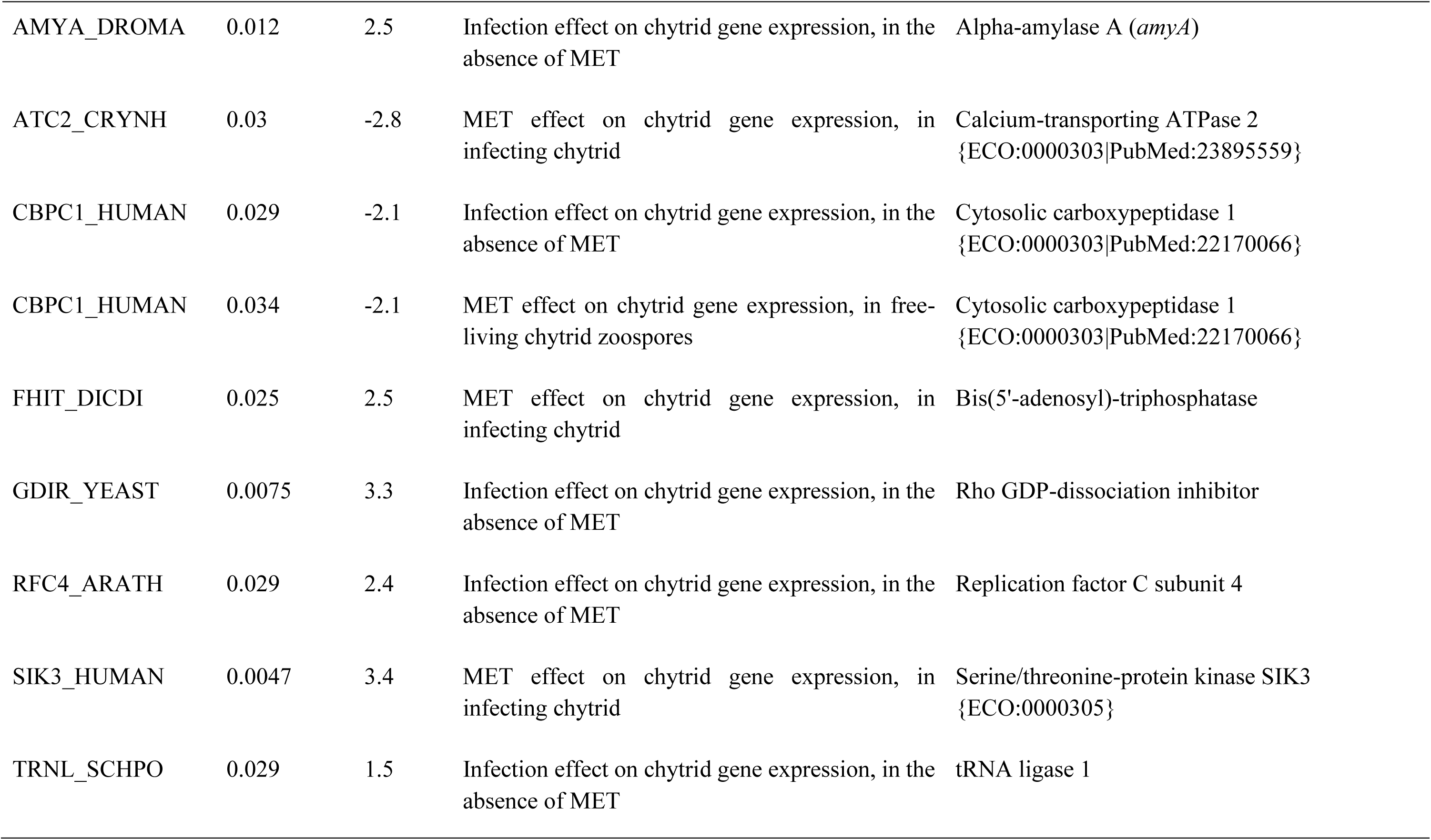
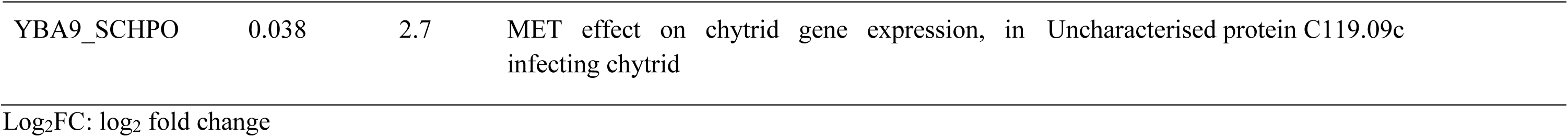
Full DEG results.

## Supplementary Figures

**Figure S1.**
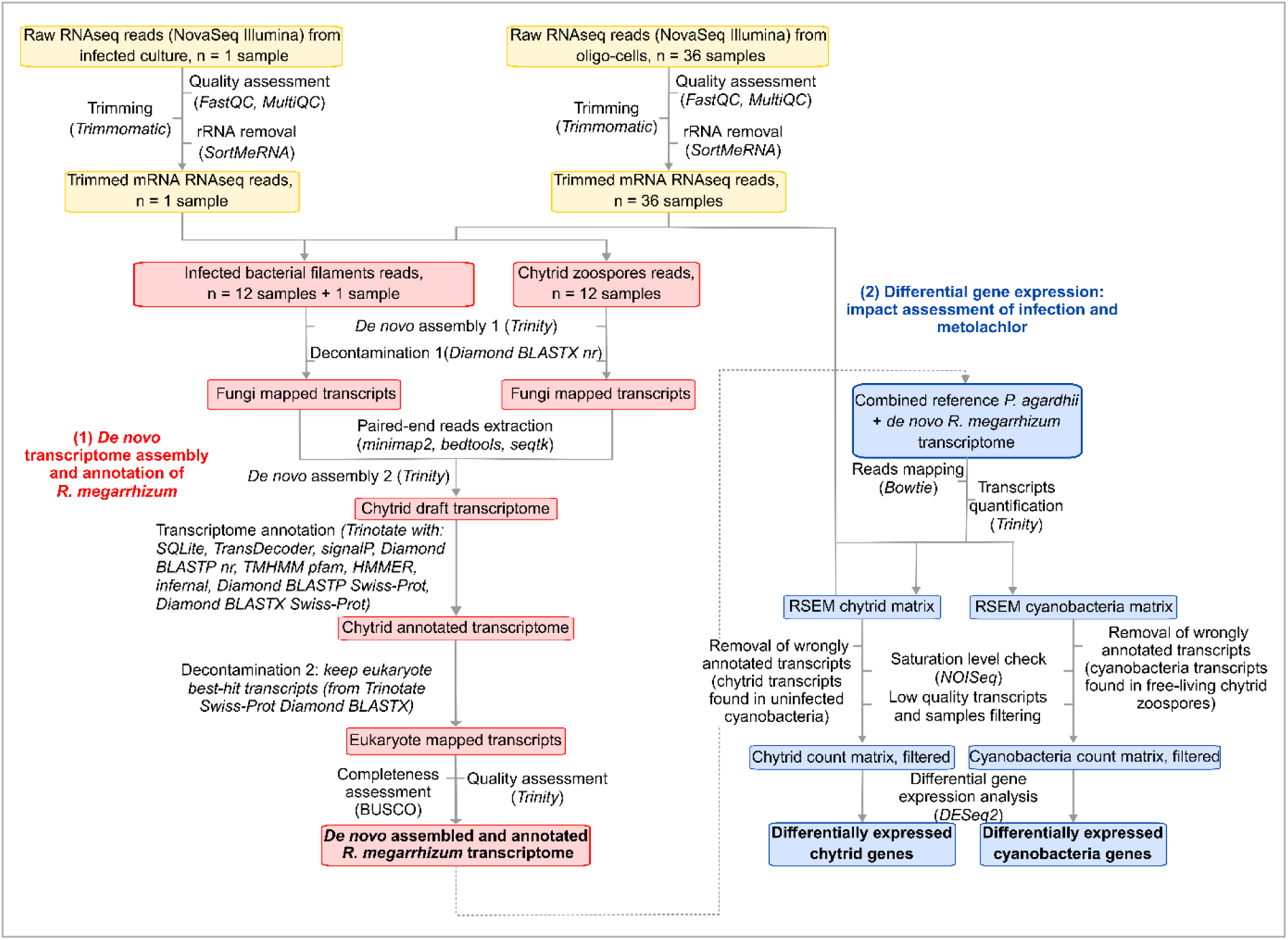
Bioinformatic pipeline.

**Figure S2.**
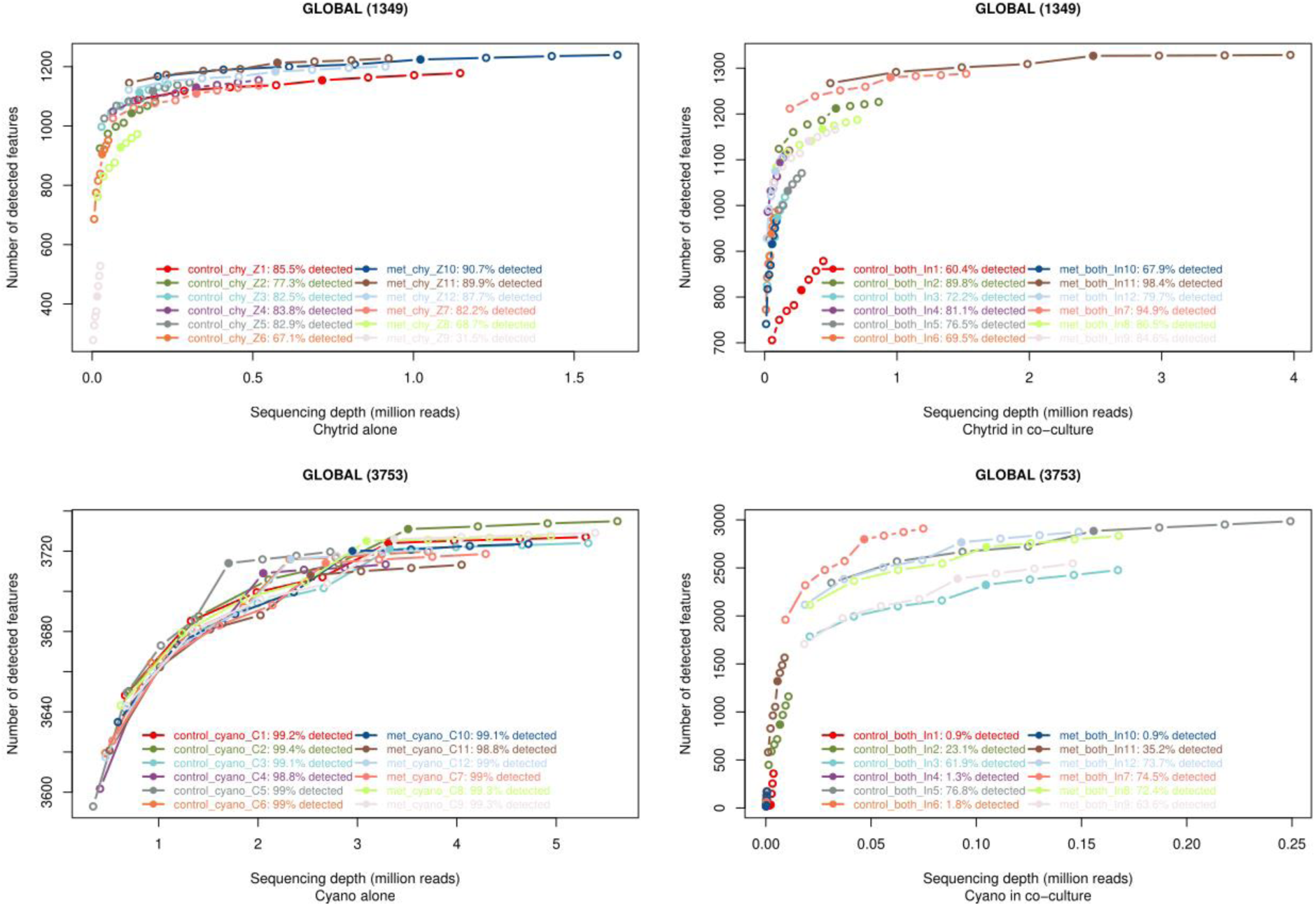
Saturation plots obtained after removing genes detected in three samples or fewer. The plots show the number of genes detected at the observed sequencing depth, as well as at simulated higher and lower depths, together with the marginal increase in gene detection per million additional reads.

**Figure S3.**
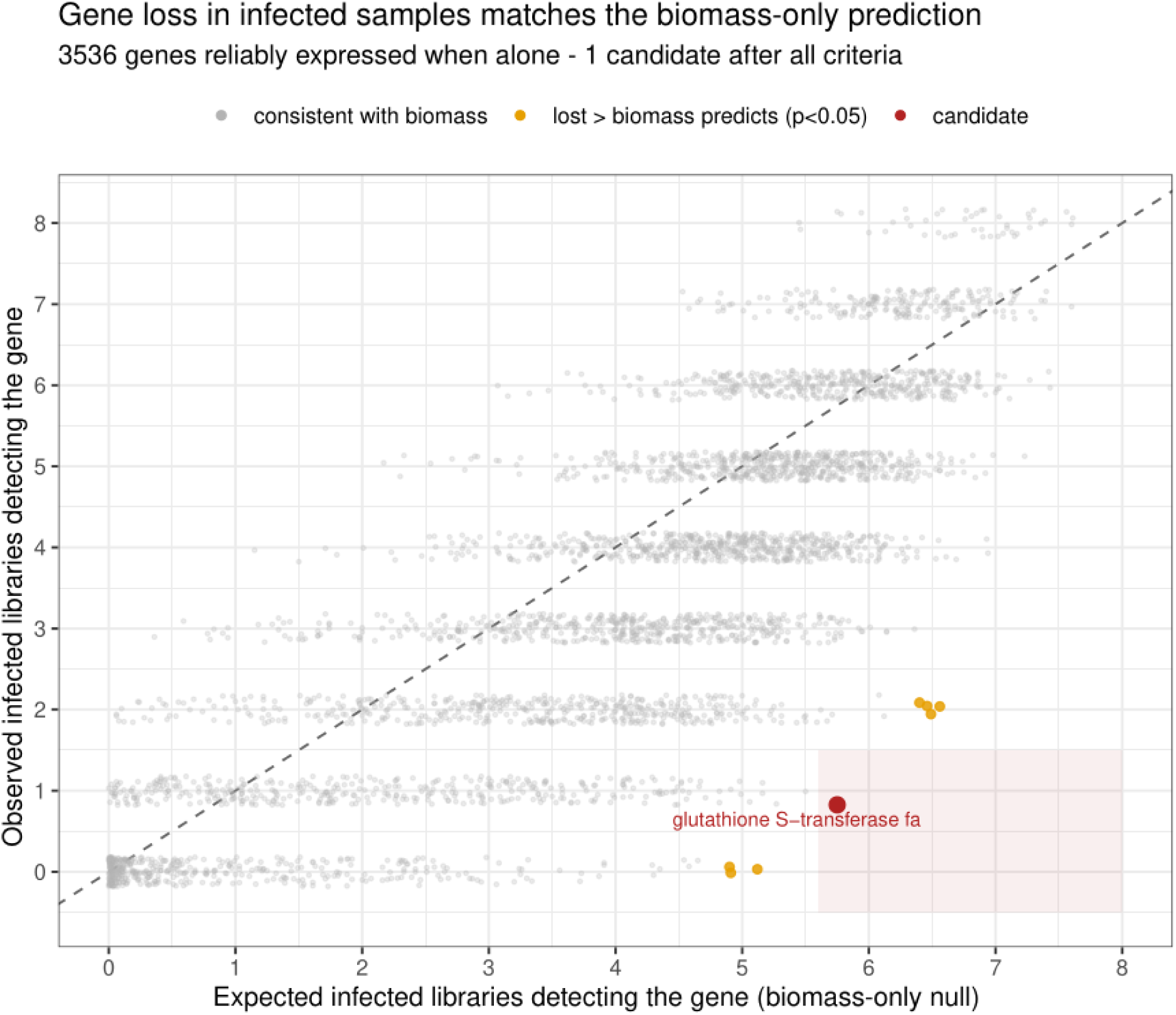
Cyanobacterial gene loss. Each point represents one cyanobacterial gene that was reliably expressed when the cyanobacterium was grown alone (≥ 5 reads in ≥ 80% of axenic libraries; *n* = 3,536 genes). The x-axis gives the number of co-culture (infected) libraries expected to detect the gene under a biomass-only null model, derived from the gene’s mean relative abundance in the uninfected transcriptome and each infected library’s cyanobacterial sequencing depth (negative binomial, dispersion = 1), and the y-axis gives the number of co-culture libraries in which it was actually detected (≥ 5 reads). The dashed line marks equality (observed = expected): genes on or above the line are detected as often as biomass alone predicts, and genes below it are detected in fewer libraries than predicted. Points are coloured grey when observed detection is consistent with the biomass-only expectation, orange when the gene was detected in significantly fewer libraries than predicted (Poisson-binomial lower-tail test, Benjamini–Hochberg adjusted *p* < 0.05), and red for the single gene that additionally met all infection-linked-loss criteria (adjusted *p* < 0.05, relative-abundance log₂ fold-change < −1, expected in ≥ 70% of infected libraries, and observed in ≤ 1); the shaded rectangle delimits this candidate region. Cyanobacterial gene loss observed during infection is likely explained by reduced biomass, while infection-specific overexpression cannot be tested.

## Notes

### Competing Interest Statement

The authors have declared no competing interest.

### Summary of Updates

This version updates the manuscript in general, improving the data analysis, readability, clarity, structure, and figures.

